# Autophosphorylation of the CK1 kinase domain regulates enzyme activity and function

**DOI:** 10.1101/660548

**Authors:** Sierra N. Cullati, Jun-Song Chen, Kathleen L. Gould

**Affiliations:** Department of Cell and Developmental Biology, Vanderbilt University Medical Center, Nashville, TN, USA

**Keywords:** CK1, casein kinase 1, kinase regulation, autoinhibition, autophosphorylation

## Abstract

CK1 enzymes are conserved, acidophilic serine/threonine kinases with a variety of critical cellular functions; misregulation of CK1 contributes to cancer, neurodegenerative diseases, and sleep phase disorders. Despite this, little is known about how CK1 activity is controlled. Here, we describe a new mechanism of CK1 autoregulation that is conserved in CK1 enzymes from yeast to human – the autophosphorylation of a threonine in the mobile L-EF loop proximal to the active site. Phosphorylation at this site inhibits kinase activity, in contrast to well-characterized T-loop autophosphorylation in other kinase families. Consequently, yeast and human enzymes with phosphoablating mutations at this site are hyperactive. In *S. pombe*, hyperactive CK1 causes defects in cell growth and morphology at a high level but protection from heat shock at a low level, highlighting the necessity of regulated CK1 function. We propose that phosphorylation on the L-EF loop prevents substrate docking with the kinase domain by shielding the positively charged binding pocket and/or sterically hindering the active site. Due to the strong sequence conservation of this autophosphorylation site and the functional importance of the L-EF loop, which is unique to the CK1 family of kinases, this mechanism is likely to regulate the majority of CK1 enzymes in vivo.

**Significance Statement:** Kinases in the CK1 family are important signaling enzymes, and they function in multiple pathways within the same cell. Misregulation of CK1 activity contributes to human disease, including cancer, neurodegenerative disease, and sleep phase disorders, yet the mechanisms that control CK1 activity are not well understood. We have identified a conserved autophosphorylation site in the CK1 kinase domain that inhibits substrate phosphorylation. We hypothesize that by using kinase domain autophosphorylation in combination with other regulatory mechanisms, CK1 enzymes can coordinate the phosphorylation of their substrates in different pathways.

## Introduction

CK1 enzymes are conserved, multifunctional kinases that are ubiquitous throughout cells and tissues (Cheong and Virshup, 2011; Knippschild et al., 2014). They regulate a variety of important cellular pathways, including mitotic checkpoint signaling (Johnson et al., 2013), DNA repair (Dhillon and Hoekstra, 1994), circadian rhythm (Eng et al., 2017; Fan et al., 2008; Meng et al., 2008; Narasimamurthy et al., 2018; Yang et al., 2017), Wnt signaling (Bernatik et al., 2011; Bryja et al., 2007; Casagolda et al., 2010; Cruciat et al., 2013; Greer and Rubin, 2011; Morgenstern et al., 2017; Peters et al., 1999; del Valle-Perez et al., 2011; Vinyoles et al., 2017), endocytosis (Peng et al., 2015; Wang et al., 2015), and neurodegenerative disease progression (Kuret et al., 2002; Li et al., 2004).

CK1 kinases have related catalytic domains (53%-98% sequence identity), plus a conserved extension to that kinase domain (KDE) that is important for enzyme stability and activity (Cheong and Virshup, 2011; Elmore et al., 2018; Greer and Rubin, 2011; Knippschild et al., 2014; Ye et al., 2016). Of the seven CK1 genes in humans, CK1δ and CK1ε share the most sequence identity with each other as well as with soluble CK1 enzymes in lower eukaryotes, including Hhp1 and Hhp2 in *Schizosaccharomyces pombe* (Dhillon and Hoekstra, 1994; Hoekstra et al., 1994). In contrast to the catalytic domains, the C-terminal tails of CK1 family members diverge in sequence and length; however, they all appear to serve as substrates of autophosphorylation. Most thoroughly studied in CK1δ, CK1ε, Hhp1, and Hhp2, autophosphorylated C-terminal tails are proposed to inhibit enzyme activity by acting as pseudosubstrates (Cegielska et al., 1998; Gietzen and Virshup, 1999; Graves and Roach, 1995; Hoekstra et al., 1994). Truncating the tail or treating the full-length enzyme with phosphatase to remove autophosphorylation was shown to increase substrate phosphorylation in vitro (Cegielska et al., 1998; Gietzen and Virshup, 1999; Graves and Roach, 1995; Hoekstra et al., 1994). In addition to C-terminal autophosphorylation, autophosphorylation of the CK1ε kinase domain has been detected (Cegielska et al., 1998), but it’s effect on CK1 activity and cellular function has never been explored.

Here, we addressed the role of kinase domain autophosphorylation in CK1δ, CK1ε, Hhp1, and Hhp2. In each case, we found that autophosphorylation of the kinase domain inhibited enzyme activity. We identified a single site of threonine autophosphorylation in CK1δ, CK1ε, and Hhp2 and two adjacent autophosphorylated threonines in Hhp1, all located on the mobile L-EF loop near the active site (Xu et al., 1995). In all four cases, phosphoablating mutations of these sites created hyperactive enzymes. Phosphorylation of these sites was also detected in Hhp1 and CK1ε purified from cells, and yeast expressing phosphoablated *hhp1* and *hhp2* were resistant to heat stress, a novel CK1 phenotype. Because the structure of the CK1 kinase domain and the location of the autophosphorylation site(s) are conserved from yeast to human, we hypothesize that kinase domain autophosphorylation works in concert with C-terminal autophosphorylation and extrinsic signaling inputs to regulate most members of the CK1 family.

## Results

### Autophosphorylation in the CK1 kinase domain inhibits activity

While autophosphorylation of full-length yeast and human CK1 enzymes is more pronounced, C-terminally truncated Hhp1, Hhp2, CK1δ, and CK1ε, (denoted by ΔC, Figure 1A) still display autophosphorylation activity, indicating that a site or sites within the kinase domain are also targeted (Figure 1B). These results are consistent with previous findings examining human CK1ε (Cegielska et al., 1998) and indicate that kinase domain autophosphorylation is a conserved feature of CK1 enzymes.

**Figure 1:**
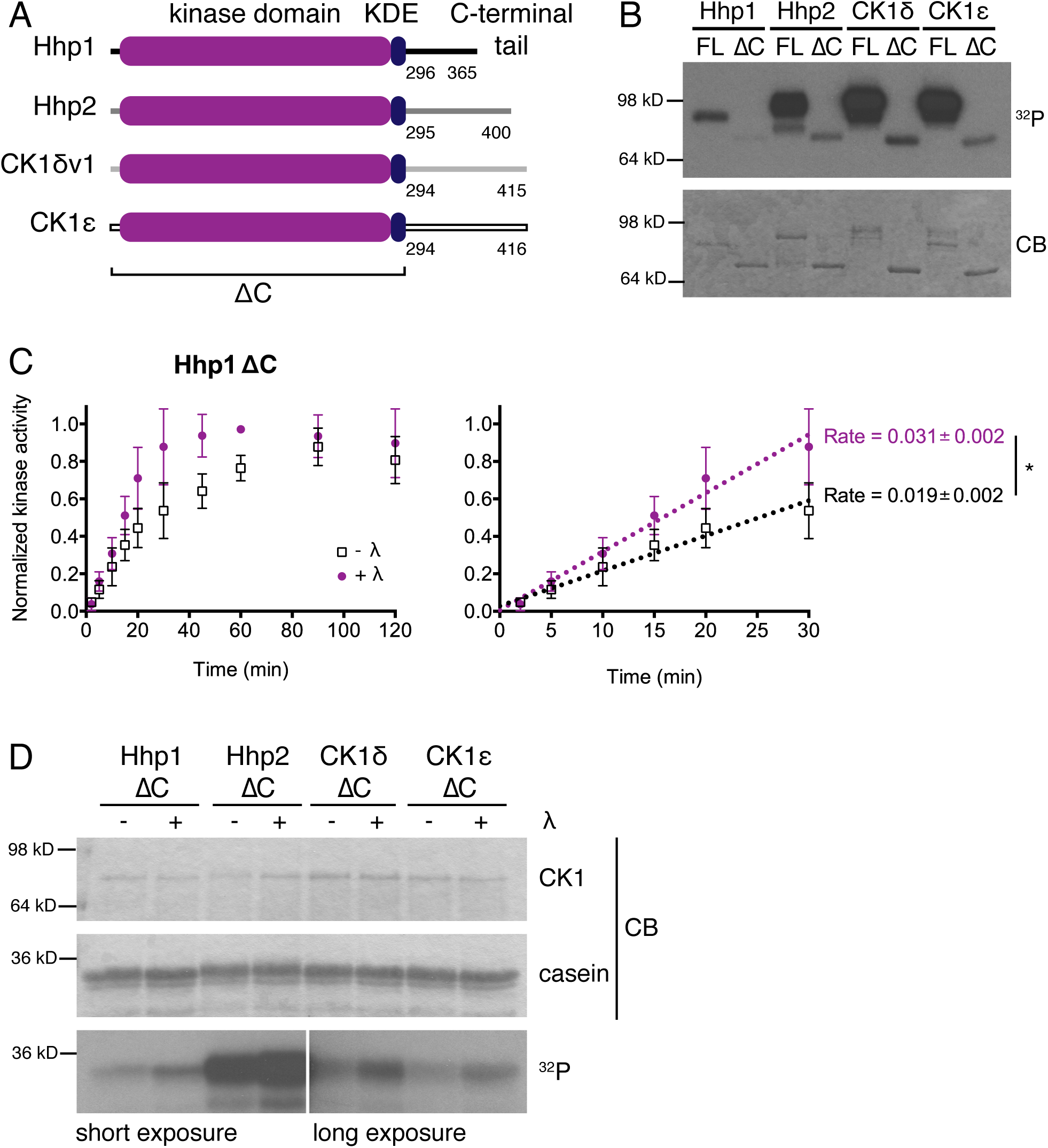
Autophosphorylation in the CK1 kinase domain inhibits activity. (A) Domain structure of CK1 enzymes used in this study. The kinase domains and kinase domain extensions (KDE) have a high degree of sequence identity from *S. pombe* to human, while the C-terminal tails are divergent. The proteins referred to as ΔC are truncated after the KDE. (B) Recombinant, MBP-tagged CK1 was incubated with γ-[^32^P]-ATP at 30°C for 30 min. Autophosphorylation was detected by autoradiography (^32^P) and total protein by Coomassie (CB). Autophosphorylation is observed even when the C-terminus is truncated. (C) MBP-Hhp1ΔC was treated with lambda phosphatase or mock treated, then incubated with casein and γ-[^32^P]-ATP at 30°C. Reactions were quenched at time points from 0-120 min, and casein phosphorylation was measured on a phosphorimager (left). Dephosphorylating the kinase domain of Hhp1 increases the rate of casein phosphorylation, as seen from the linear segment of the curve (right). Data from three independent replicates is shown as the mean ± SD. The rate is equal to the slope of the regression line that was calculated in Prism. (D) Truncated, MBP-tagged CK1 was treated with lambda phosphatase or mock treated, then incubated with casein and γ-[^32^P]-ATP at 30°C for 45 min. Casein phosphorylation was detected by autoradiography (^32^P) and total protein by Coomassie (CB). Dephosphorylation of the kinase domain increases the activity of all four enzymes toward substrate.

We and others have found that these enzymes autophosphorylate when expressed in bacteria (Cegielska et al., 1998; Gietzen and Virshup, 1999; Graves and Roach, 1995; Hoekstra et al., 1994), and interestingly, pretreating recombinant Hhp1ΔC with lambda phosphatase increased kinase activity (Figure 1C). We performed a kinetics experiment to measure the phosphorylation of casein by Hhp1ΔC in the presence of γ-[^32^P]-ATP, and we found that phosphatase treatment of the enzyme increased the reaction rate ∼1.6 fold (Figure 1C). PP2A also activated CK1εΔC about 2 fold toward SV40 T-antigen (Cegielska et al., 1998). Furthermore, endpoint in vitro kinase assays with Hhp1ΔC, Hhp2ΔC, CK1δΔC, and CK1εΔC all showed that dephosphorylation of CK1 increased the amount of ^32^P incorporated into casein after 45 minutes (Figure 1D). These data suggested that kinase domain autophosphorylation might inhibit kinase activity similarly to what has been reported for C-terminal autophosphorylation (Cegielska et al., 1998; Gietzen and Virshup, 1999; Graves and Roach, 1995; Hoekstra et al., 1994).

### CK1 enzymes autophosphorylate a conserved threonine in the kinase domain

To test this idea more completely, we determined the site(s) of kinase domain autophosphorylation. Phosphoamino-acid analysis revealed that autophosphorylation in Hhp1ΔC, Hhp2ΔC, CK1δΔC, and CK1εΔC occurs exclusively on threonine (Supplemental Figure 1A). While a previous study proposed that Hhp1 and Hhp2 were dual-specificity kinases capable of tyrosine phosphorylation (Hoekstra et al., 1994), we did not detect tyrosine autophosphorylation on yeast or human CK1 enzymes. To identify the phosphorylated threonine(s), we used *S. pombe* Hhp1 as a model and analyzed tandem affinity purifications of *hhp1* tagged at the endogenous locus (Supplemental Figure 1B), as well as recombinant full-length Hhp1 and Hhp1ΔC by mass spectrometry (Supplemental Figure 1C). In all samples, the most abundant phosphopeptides that we identified contained phosphorylated T221. Further phosphopeptide mapping of in vitro phosphorylated Hhp1ΔC wildtype (WT), T221V, T222V, and T221V/T22V (henceforth called VV) confirmed that both T221 and the adjacent T222 are sites of autophosphorylation (Supplemental Figure 2A).

An alignment of CK1 sequences indicated that T222 is conserved in all other CK1 enzymes examined from yeast to human CK1 (Figure 2A). Thus, we tested if the homologous residues (T221 in *S. pombe* Hhp2, T220 in human CK1δ and CK1ε) were autophosphorylated. We identified Hhp2 phospho-T221 and CK1δ phospho-T220 by mass spectrometry of recombinant protein, and CK1ε phospho-T220 in multifunctional tandem affinity purifications from HEK293 cells (Supplemental Figure 1D-F). These residues were confirmed as autophosphorylation sites by phosphopeptide mapping (Supplemental Figure 2B-D). Consistent with the idea that kinase domain autophosphorylation is a conserved feature of CK1 enzymes, mutation from threonine to asparagine abolished kinase domain autophosphorylation in each case (Figure 2B).

**Figure 2:**
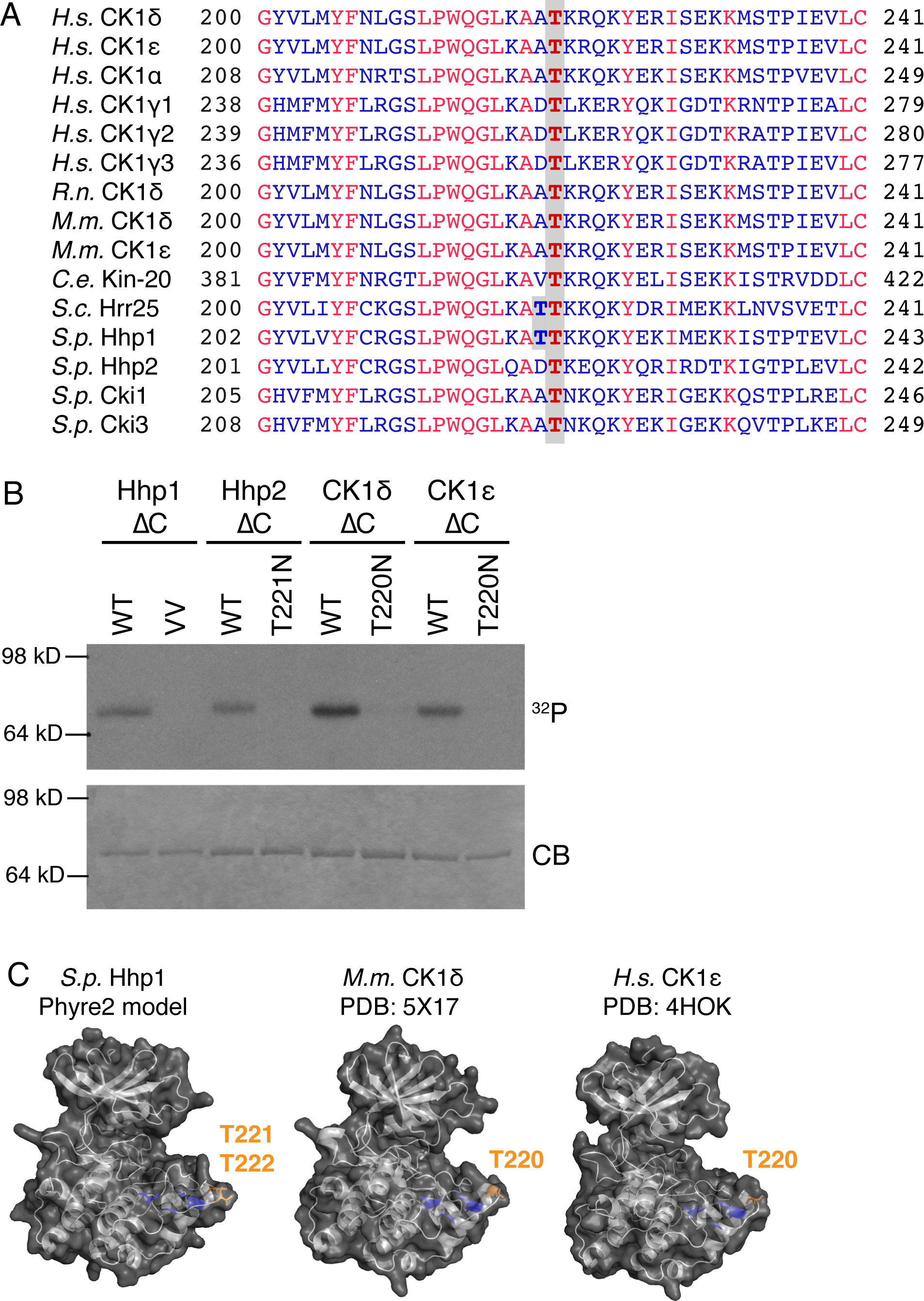
CK1 autophosphorylates a conserved threonine in the kinase domain. (A) CK1 enzymes from multiple species were aligned using COBALT. Amino acids in red were conserved in all proteins; those in blue had one or more conservative substitutions. The threonines that align with kinase domain autophosphorylation sites in Hhp1 are boxed in grey. *H.s., Homo sapiens; R.n., Rattus norvegicus; M.m., Mus musculus; C.e. Caenorhabditis elegans; S.c. Saccharomyces cerevisiae; S.p. Schizosaccharomyces pombe*. (B) Truncated, MBP-tagged CK1 was incubated with γ-[^32^P]-ATP at 30°C for 30 min. Autophosphorylation was detected by autoradiography (^32^P) and total protein by Coomassie (CB). Mutating the conserved kinase domain threonine abolishes autophosphorylation. (C) Homology model of yeast Hhp1ΔC generated using the Phyre2 server (Kelley et al., 2015) alongside crystal structures of mouse CK1δ (Shinohara et al., 2017) and human CK1ε (Long et al., 2012). Kinase domain autophosphorylation sites are in orange, and a basic patch hypothesized to interact with substrates (Longenecker et al., 1996; Xu et al., 1995; Ye et al., 2016) is in blue.

Interestingly, mutation to alanine in any of the four enzymes tested or to valine in Hhp2, CK1δ, and CK1ε not only abolished autophosphorylation but also disrupted kinase activity towards exogenous substrate, suggesting that this conserved region of the protein is highly sensitive to perturbation (Supplemental Figure 3A-C). A homology model of *S. pombe* Hhp1 generated by the Phyre2 toolkit (Kelley et al., 2015) localizes the autophosphorylated threonines to a loop proximal to the active site (Figure 2C, left), defined as L-EF in the first CK1 structure (Xu et al., 1995). Electron density corresponding to this region of the protein is not defined in most CK1 structures, including *S. pombe* Cki1 (Mashhoon et al., 2000; Xu et al., 1995) and rodent CK1δ (Longenecker et al., 1996, 1998; Zeringo et al., 2013). The two structures with ordered density in this loop (Figure 2C, Long et al., 2012; Shinohara et al., 2017) show T220 from mouse CK1δ and human CK1ε in a similar position as T221/T222 in the Hhp1 model, and again, the L-EF loop has large B-factors and was highly fluctuated in molecular dynamics simulations (Shinohara et al., 2017). This suggests that these sites reside in a minimally structured, mobile loop and that the position and/or dynamics of L-EF may be important for CK1 function.

### Ablating CK1 autophosphorylation promotes substrate phosphorylation

Since dephosphorylation of C-terminally truncated CK1 increased the ability to phosphorylate casein (Figure 1C-D), we tested whether mutations that abolish autophosphorylation at this site also increased the activity of Hhp1, Hhp2, CK1δ, and CK1ε. All four kinases displayed the same pattern of activation, though the basal level of activity and the extent to which mutation or dephosphorylation increased activity depended on the protein. As we saw for the truncated enzymes (Figure 1D), lambda phosphatase treatment of full-length Hhp1, Hhp2, CK1δ, and CK1ε increased their activity toward exogenous substrate in endpoint assays (Figure 3A-D). When we mutated the kinase domain threonine, all eight mutants phosphorylated casein to a greater extent than the corresponding WT enzymes (Figure 3A-D). This indicates that even in the full-length enzymes that have autophosphorylated their C-termini, abolishing autophosphorylation in the kinase domain increases activity. Furthermore, in C-terminally truncated CK1s, the threonine mutations increased kinase activity to the same extent as lambda phosphatase treatment of the WT enzyme, and phosphatase treatment of the threonine mutants did not lead to an additional increase in kinase activity (Figure 3A-D and Figure 4C).

**Figure 3:**
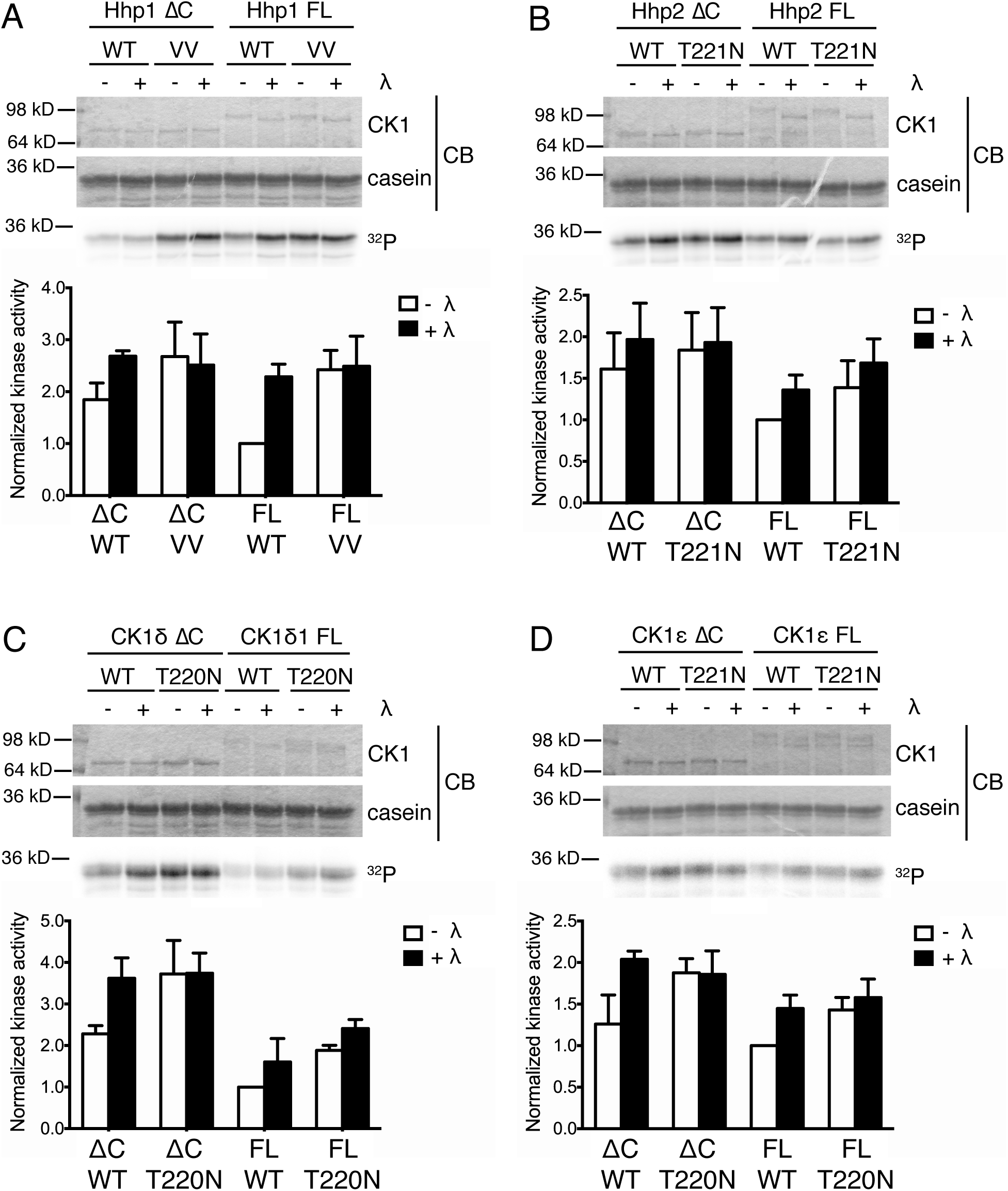
Ablation of kinase domain autophosphorylation increases CK1 activity in vitro. In both full-length (FL) and truncated (ΔC) Hhp1 (A), Hhp2 (B), CK1δ (C) and CK1ε (D), mutation of the conserved kinase domain threonine(s) increases phosphorylation of casein. MBP-tagged CK1 enzymes were treated with lambda phosphatase or mock treated, then incubated with casein and γ-[^32^P]-ATP at 30°C for 45 min. Phosphorylated casein was quantified on a phosphorimager (^32^P), and total protein was visualized by Coomassie (CB). Bar graphs represent the mean ± SD of 3-4 independent replicates, which have been normalized to each FL, WT enzyme.

**Figure 4:**
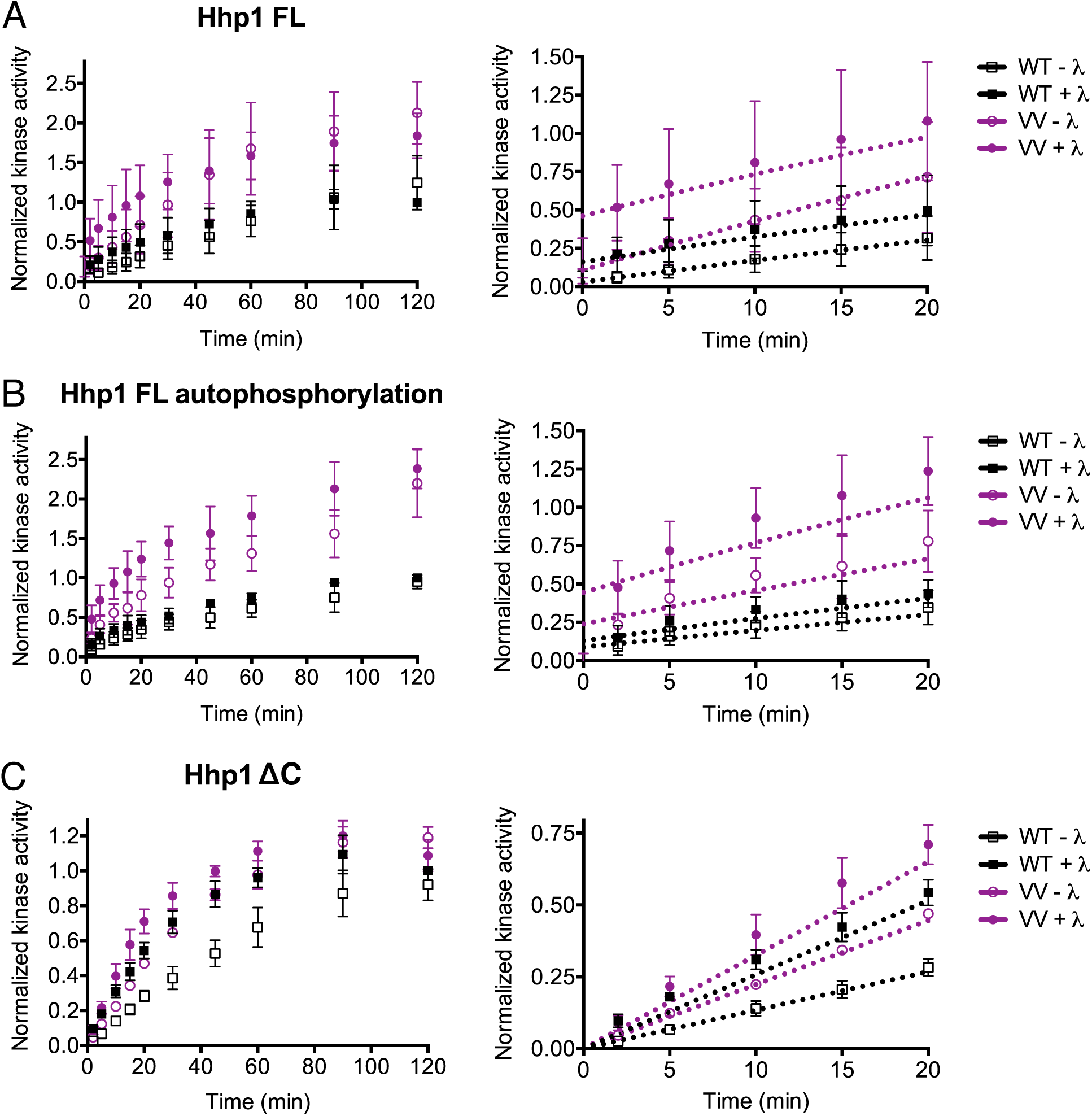
Autophosphorylation of Hhp1 T221/T222 decreases the rate of substrate phosphorylation and C-terminal autophosphorylation. The indicated MBP-Hhp1 enzymes were treated with lambda phosphatase or mock treated, then incubated with casein and γ-[^32^P]-ATP at 30°C. Reactions were quenched at time points from 0-120 min, and casein phosphorylation (A and C) or autophosphorylation (B) was measured on a phosphorimager. The full timecourse is on the left of each panel, while the linear section of the curve is enlarged on the right. FL Hhp1 VV has increased activity toward casein, as well as increased autophosphorylation of C-terminal sites. Truncated Hhp1-VV phosphorylated casein to the same extent as lambda phosphatase-treated WT and was not further activated by phosphatase treatment.

To better quantify the increase in kinase activity that results from preventing autophosphorylation of the kinase domain, we used Hhp1 as a model CK1 enzyme and performed kinetics experiments to measure the phosphorylation of casein over time in the presence of γ-[^32^P]-ATP (Figure 4A and C). At saturation, full-length Hhp1 VV was approximately twice as active as Hhp1 WT toward casein (Figure 4A) and also displayed an increase in autophosphorylation activity that presumably reflects increased phosphorylation of the C-terminus (Figure 4B). For Hhp1ΔC, the rate of casein phosphorylation for the phosphatase-treated WT enzyme and the VV mutant plus and minus phosphatase were nearly identical to each other, and all were greater than the autophosphorylated enzyme (Figure 4C). Taken together, these results indicate that the phosphorylation status of the kinase domain threonine significantly affects CK1 activity in vitro.

### Ablating CK1 autophosphorylation produces gain-of-function alleles in vivo

Deletion of *hhp1* and *hhp2* is known to cause several cellular phenotypes in yeast, including delayed repair of DNA double strand breaks (Dhillon and Hoekstra, 1994) that increases sensitivity to DNA damage (Bimbó et al., 2005; Chen et al., 2015; Dhillon and Hoekstra, 1994), defects in meiosis (Ishiguro et al., 2010; Phadnis et al., 2015), increased cell length (Dhillon and Hoekstra, 1994), and increased monopolar growth (Koyano et al., 2010). However, the effects of overexpression or gain-of-function are poorly characterized. We found that overexpressing *hhp1, hhp1-VV*, and the kinase-dead *hhp1-K40R* from a plasmid under the control of the *nmt1* promoter (Basi et al., 1993; Forsburg, 1993) caused cell death, while overexpressing *hhp2, hhp2-T221N*, and *hhp2-K41R* caused a milder growth defect (Figure 5A). The morphology of *hhp1-K40R* and *hhp2-K41R* cells was similar to that of *hhp1Δ*, namely that cells were long (Dhillon and Hoekstra, 1994), and we hypothesize that these kinase-dead alleles act as dominant negatives (Supplemental Figure 4A). In contrast, *hhp1, hhp1-VV, hhp2,* and *hhp2-T221N* displayed polarity defects – branched cells and round cells (Supplemental Figure 4A) – as well as multiseptated cells and cut phenotypes, which indicate problems in cytokinesis (Figure 5B). The similarity between the phenotypes of the autophosphorylation mutants and the wildtype strains suggests that, consistent with our in vitro data, these mutations increase kinase activity in cells, thereby behaving as gain-of-function alleles.

**Figure 5:**
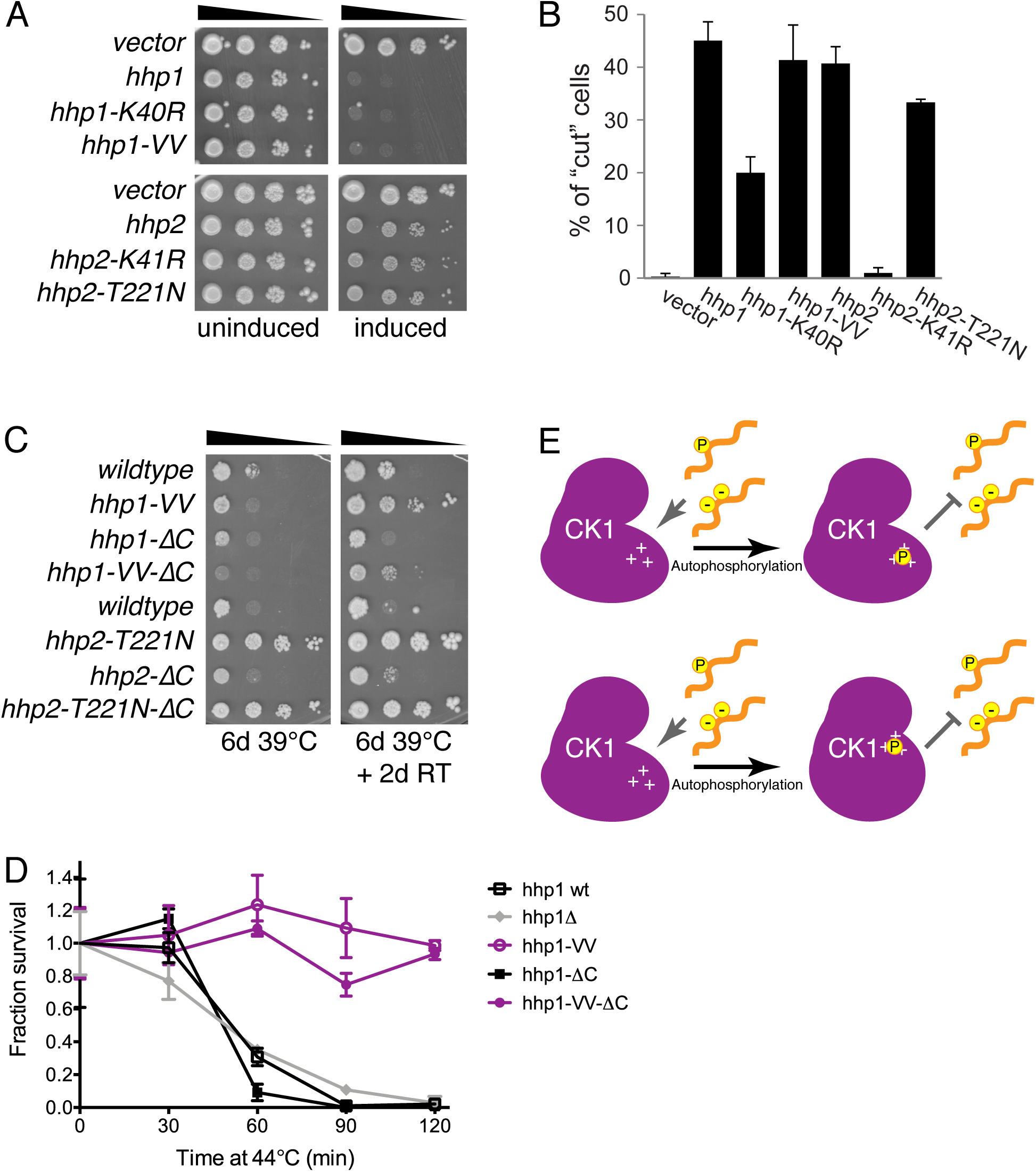
Ablation of kinase domain autophosphorylation causes gain-of-function phenotypes in vivo. (A) The indicated *pREP1-hhp1* and *pREP1-hhp2* plasmids were transfected into *S. pombe*, and expression was induced by thiamine washout. Ten-fold serial dilutions were spotted on plates with (uninduced) and without (induced) thiamine and incubated at 32°C. (B) Overexpression of the indicated alleles was induced by thiamine washout. Cells were imaged at 25°C, and the percent of the population exhibiting a cut phenotype (septation despite abnormal nuclear division) was quantified in n = 100 cells in each of 3 replicates. Bars indicate mean ± SD. (C) The indicated alleles were integrated at the endogenous loci, and 10-fold serial dilutions were spotted. *hhp2-T221N* strains grew better than *wildtype* at 39°C (left), and *hhp1-VV* strains were able to recover from heat stress better than *wildtype* (right). (D) Viability of the indicated strains after 0-120 minutes of heat shock. Strains were grown to log-phase at 29°C, transferred to 44°C, and aliquots were diluted and plated in duplicate every 30 minutes. Colonies were counted after 3d at 29°C. (E) Kinase domain autophosphorylation may inhibit CK1 activity by preventing interaction with substrate. We hypothesize that kinase domain autophosphorylation may mask the positive charge of the substrate binding patch (top) and/or cause a conformational change that occludes substrates from the active site (bottom).

We next determined the effects of ablating kinase domain autophosphorylation at physiological expression levels. We constructed strains of *S. pombe* with *hhp1* and *hhp2* mutants integrated at their endogenous loci and measured cell growth at 19-36°C (Supplemental Figure 4B). Truncating the C-terminus of *hhp1* or *hhp2* does not affect cell growth at these temperatures (Elmore et al., 2018), and mutating the kinase domain threonines in Hhp1 and Hhp2 increased kinase activity to a similar level (Figure 3A-B), so we hypothesized that cells may be buffered to these relatively small (∼1.5-3 fold) increases in CK1 activity. Indeed, strains harboring *hhp1-VV, hhp1-VV-ΔC, hhp2-T221N,* and *hhp2-T221N-ΔC* grew normally at these temperatures (Supplemental Figure 4B). Surprisingly, we found that at a high temperature (39°C) *hhp2-T221N* and *hhp2-T221N-ΔC* grew better than *wildtype*, and *hhp1-VV* and *hhp1 VV-ΔC* grew better than *wildtype* when the same plates were allowed to recover from heat shock at room temperature (Figure 5C). This indicates that CK1 gain-of-function may confer heat resistance to *S. pombe*, a phenotype that has not previously been described. To quantify recovery from heat shock at different levels of CK1 activity, we grew *hhp1 wt* and *hhp1-VV* strains to log phase at 29°C, then transferred to 44°C. Aliquots were removed at 30-minute intervals and plated to score survival at 29°C (Figure 5D). *hhp1 wt, hhp1-ΔC,* and *hhp1Δ* cells lost most of their viability after 60 minutes of heat shock and were completely inviable after 120 minutes. *hhp1-VV* and *hhp1-VV-ΔC*, however, did not lose viability even after two hours at 44°C. These data show that gain-of-function mutations that cause an increase in CK1 activity can be protective at a low level but detrimental to the organism at a high level. Therefore, regulatory mechanisms must buffer CK1 activity to within a reasonable range and/or to modulate the phosphorylation of CK1 substrates in vivo.

## Discussion

### Significance of autophosphorylation in the CK1 kinase domain

Although CK1 enzymes are multifunctional kinases involved in a variety of essential signaling pathways related to human disease, the prevailing idea has been that they are not regulated – either CK1s are constitutively active (Knippschild et al., 2005), or they are constitutively inactivated by C-terminal autophosphorylation until a phosphatase releases them from the inhibited state (Rivers et al., 1998). This dichotomy raises the question of to what extent and by which mechanisms CK1 activity is controlled, especially within the context of a living cell, but answering this question has been challenging. The primary focus of previous research has been on extrinsic regulation of CK1 substrates by phosphatases and priming kinases, or on C-terminal autophosphorylation (Bhandari et al., 2013; Cegielska et al., 1998; Gietzen and Virshup, 1999; Graves and Roach, 1995; Hoekstra et al., 1994; Ishiguro et al., 2010; Rivers et al., 1998; Zeng et al., 2005). More recent work has also shown that CK1 activity can be modulated by binding interactions and subcellular targeting (Cruciat et al., 2013; Elmore et al., 2018; Greer and Rubin, 2011). Here we show for the first time that autophosphorylation of the CK1 kinase domain, independently of the C-terminus or of other kinases and phosphatases, inhibits enzyme activity.

In general, the kinase domains of CK1 enzymes are emerging as more important arbiters of function than has been previously appreciated. Residues at the base of the kinase domain mediate subcellular localization in order to compartmentalize CK1 signaling at the spindle pole (Elmore et al., 2018). The kinase domains of fly and mammalian CK1 are involved in stable substrate interactions with PERIOD proteins (Dahlberg et al., 2009; Lee et al., 2009; Preuss et al., 2004; Vielhaber et al., 2000). The CK1δ and CK1ε kinase domains alone display temperature compensated phosphorylation of circadian substrates (Isojima et al., 2009; Qin et al., 2015; Shinohara et al., 2017; Zhou et al., 2015), and in fact, the presence of the phosphorylated C-terminus appears to increase the Q_10_, making the temperature compensation less robust (Isojima et al., 2009). Given that the model of autoinhibition mediated by the C-terminus requires an interaction with the kinase domain, it would be interesting to determine whether autophosphorylation in the kinase domain affects this interaction and thereby the occupancy of C-terminal autophosphorylation sites. Lastly, while much of the focus has been on CK1 C-termini, it is the kinase domain that is conserved, so mechanisms of regulation intrinsic to the kinase domain may be ones selected by evolutionary pressure.

### Potential mechanisms of kinase domain-mediated autoinhibition

We found that kinase activity is highly sensitive to mutagenesis in the conserved L-EF loop where the kinase domain autophosphorylation site is located (Supplemental Figure 3), suggesting that the position and/or dynamics of L-EF is important for CK1 function. The residues of L-EF comprise an insertion unique to the CK1 family that may participate in anion binding (Longenecker et al., 1996; Xu et al., 1995); this loop was also found to confer temperature compensation to the phosphorylation rate of substrates involved in circadian rhythms (Shinohara et al., 2017). In other CK1 structures, anions have been found to bind a conserved basic patch near the active site (Longenecker et al., 1996; Xu et al., 1995; Ye et al., 2016), leading to the hypothesis that this site may interact with negatively charged substrates in vivo. Given this previous work and our current findings, we propose that threonine autophosphorylation may regulate the movement of the L-EF loop such that the negative phosphate group interacts with the basic substrate recognition patch. This could shield the positive charge that recruits substrates (Figure 5E, top) and/or cause a conformational change that occludes substrate access to the active site (Figure 5E, bottom), thus providing a mechanism for autoinhibition. A structure of the active enzyme, perhaps in complex with substrate, would shed light on the molecular details by which kinase domain autophosphorylation inhibits substrate phosphorylation.

This inhibitory mechanism is in sharp contrast to the more common paradigm of activating T-loop phosphorylation. Many serine/threonine and all tyrosine kinases have an arginine directly upstream of the catalytic aspartate, and these RD kinases often utilize T-loop phosphorylation to properly align the catalytic aspartate (Beenstock et al., 2016; Johnson et al., 1996). The L-9D loop of CK1 is homologous to the T-loop, though the terminal APE motif is replaced with SIN, and while CK1 enzymes are RD kinases, they do not require T-loop phosphorylation for activity (Beenstock et al., 2016; Johnson et al., 1996). The autophosphorylated threonine residue(s) on L-EF seem to gate the substrate binding channel rather than position the active site residues. A second conserved basic patch in CK1 appears to overlap with the T-loop phosphorylation site of other kinases, leading to the hypothesis that it may interact with autophosphorylated residues in the kinase domain (Xu et al., 1995) or the C-terminus (Longenecker et al., 1996; Ye et al., 2016). Anion binding in this site may alter the conformation of L-9D to decrease kinase activity (Ye et al., 2016), but this proposed mechanism is distinct from the one described here; the activating effect of phosphoablating mutations even in full-length, C-terminally phosphorylated CK1 (Figure 3, Figure 4A) argues that they are separate. Mutating threonines in the L-9D loop of Hhp1 did not change the enzyme’s autophosphorylation (Supplemental Figure 2E), confirming that these sites are not autophosphorylated, though they could still be targeted by other kinases in vivo.

### Physiological consequences of CK1 misregulation

Dramatically overexpressing *hhp1* or *hhp2* in *S. pombe* impairs cell viability (Figure 5A), but low-level activation achieved by ablating kinase domain autophosphorylation promotes cell survival (Figure 5C-D). The ability of *hhp1-VV* and *hhp2-T221N* to protect against heat shock suggests a novel function of yeast CK1 in heat resistance. This function, as well as the many others ascribed to CK1, clearly must be regulated in cells to ensure that the right cohort of substrates is phosphorylated to the right extent under the right circumstances. There is evidence that PP1 (Rivers et al., 1998), PP2A (Rivers et al., 1998), calcineurin (Liu et al., 2002), and PP5 (Partch et al., 2006) can dephosphorylate CK1ε to increase activity, though this was presumed to occur on C-terminal autophosphorylation sites. Whether the kinase domain threonine is dephosphorylated, under what conditions, and by which phosphatase(s) is unclear. Furthermore, the interplay between kinase domain and C-terminal autophosphorylation, as well as other mechanisms of CK1 regulation such as allosteric conformational changes induced by binding partners (Cruciat et al., 2013) and phosphorylation by other kinases (Bischof et al., 2013; Giamas et al., 2007; Ianes et al., 2016; Meng et al., 2016), remains to be elucidated.

## Materials and Methods

### Molecular biology and protein purification

All plasmids were generated by standard molecular biology techniques. *hhp1* and *hhp2* mutants were created by mutagenizing pIRT2 plasmids containing *hhp1*^*+*^ and *hhp2*^*+*^ using a QuikChange site-directed mutagenesis kit (Agilent Technologies). For protein production, cDNA for Hhp1, Hhp2, CK1δv1, and CK1ε was mutagenized in the pMAL-C2 vector. CK1 ΔC constructs consisted of the following amino acids: Hhp1 1-296, Hhp2 1-295, CK1δ 1-296, CK1ε 1-294. For overexpression, *hhp1*^*+*^ and *hhp2*^*+*^ were cloned into pREP1, then mutagenized. Plasmids were validated by DNA sequencing.

Protein production was induced in *Escherichia coli* Rosetta2(DE3)pLysS cells by addition of 0.4mM IPTG overnight at 17°C. Cells were lysed using 300µg/mL lysozyme for 20 minutes followed by sonication. MBP fusion proteins were purified on amylose beads (New England Biolabs) in column buffer (20mM Tris pH 7.4, 150mM NaCl, 1mM EDTA, 0.1% NP40, 1mM DTT, 1mM PMSF, 1.3mM benzamidine, protease inhibitor tablets [Roche]) and eluted with maltose (20mM Tris pH 7.4, 150mM NaCl, 1mM EDTA, 1mM DTT, 1mM PMSF, 1.3mM benzamidine, 10mM maltose, 10% glycerol).

### In vitro kinase assays

Kinases were treated with 0.5µL lambda phosphatase (New England Biolabs) per 1µg protein for 1 hour at 30°C in PMP buffer (New England Biolabs) plus 1mM MnCl_2_. For negative controls, an equivalent volume of buffer was added instead of phosphatase. Phosphatase reactions were quenched by the addition of 8mM Na_3_VO_4_ immediately prior to the kinase assay.

Autophosphorylation reactions were performed with 500ng kinase in PMP buffer plus 100μM cold ATP, 0.5µCi γ-[^32^P]-ATP, and 10mM MgCl_2_ at 30°C for 30 min. Reactions were quenched by boiling in SDS-PAGE sample buffer and proteins were separated by SDS-PAGE. Phosphorylated proteins were visualized by autoradiography and total protein by Coomassie blue staining.

Casein phosphorylation reactions were performed with 300ng kinase and 25µM dephosphorylated alpha casein (Sigma) in PMP buffer plus 100μM cold ATP, 1µCi γ-[^32^P]-ATP, and 10mM MgCl_2_ at 30°C for 45 min. Reactions were quenched by boiling in SDS-PAGE sample buffer. Kinetic assays were performed using 200ng kinase and 25µM casein in PMP buffer plus 250μM cold ATP, 1µCi γ-[^32^P]-ATP, and 10mM MgCl_2_. Reactions were quenched at time points from 0-120 min by boiling in SDS-PAGE sample buffer. Proteins were separated by SDS-PAGE and stained in Coomassie, then gels were dried. Phosphorylated proteins were visualized using an FLA7000IP Typhoon Storage Phosphorimager (GE Healthcare Life Sciences) and quantified in ImageJ. Relative kinase activity was plotted in Prism.

### Mass spectrometry methods

TCA-precipitated proteins from in vitro kinase assays or tandem-affinity purifications were subjected to mass spectrometric analysis on an LTQ Velos (Thermo) by 3-phase multidimensional protein identification technology (MudPIT) as previously described (Chen et al., 2013) with the following modifications. Proteins were resuspended in 8M urea buffer (8M urea in 100 mM Tris, pH 8.5), reduced with Tris (2-carboxyethyl) phosphine, alkylated with 2-chloro acetamide, and digested with trypsin or elastase. The resulting peptides were desalted by C-18 spin column (Pierce). For the kinase assay samples 6 salt elution steps were used (i.e. 25, 50, 100, 600, 1000, and 5000 mM ammonium acetate) instead of the full 12 steps for TAP samples. Raw mass spectrometry data were filtered with Scansifter and searched by SEQUEST algorithm. Scaffold (version 3.6.0 or version 4.2.1) and Scaffold PTM (version 3.0.1) (both from Proteome Software, Portland, OR) were used for data assembly and filtering. The following filtering criteria were used: minimum of 90.0% peptide identification probability, minimum of 99% protein identification probability, and minimum of two unique peptides. TAP (Johnson et al., 2013) and MAP (Ma et al., 2012) purifications were done as described previously.

### Phospho-amino acid analysis and phosphopeptide mapping

Autophosphorylation reactions were performed with 4µg lambda phosphatase-treated kinase in PMP buffer plus 100μM cold ATP, 4µCi γ-[^32^P]-ATP, and 10mM MgCl_2_ at 30°C for 30 min. Reactions were quenched by boiling in SDS-PAGE sample buffer and proteins were separated by SDS-PAGE. Phosphorylated proteins were transferred to PVDF membranes. For PAA, proteins were subjected to partial acid hydrolysis using boiling 6M HCl for 1 hr. Hydrolyzed amino acids were separated by two-dimensional thin-layer electrophoresis (Boyle et al., 1991). For phosphopeptide mapping, proteins were digested off the membrane with 10µg trypsin at 37°C overnight. Peptides were lyophilized and resuspended in pH 1.9 buffer. Tryptic peptides were separated in the first dimension by thin-layer electrophoresis and in the second dimension by chromatography (Boyle et al., 1991). After separations, TLC plates were exposed to film for 2-4 d at −80°C with intensifying screens.

### Yeast methods

*S. pombe* strains used in this study (Supplemental Table 1) were grown in yeast extract (YE) media (Moreno et al., 1991). Crosses were performed in glutamate medium (Moreno et al., 1991), and strains were constructed by tetrad analysis. For *hhp1* and *hhp2* gene replacements, haploid *hhp1::ura4+* and *hhp2::ura4+* strains were transformed using standard lithium acetate methods (Keeney and Boeke, 1994) to introduce linear *hhp1-VV* and *hhp2-T221N* gene fragments generated by digestion of pIRT2-hhp1-VV and pIRT2-hhp2-T221N plasmids with BamHI and PstI. Integrants were selected based on resistance to 1.5mg/ml 5-fluoroorotic acid (Fisher Scientific) and validated by whole-cell PCR using primers homologous to endogenous flanking sequences in combination with those within the ORF. Truncations were generated by insertion of the *kanMX6* or *natMX6* cassette from pFA6 as previously described (Bähler et al., 1998), followed by selection on G418 (100µg/mL; Sigma-Aldrich, St. Louis, MO) or nourseothricin (100µg/mL; Goldbio, St. Louis, MO). Insertions were validated by whole-cell PCR using primers homologous to the resistance cassette and the endogenous ORF. All constructs and integrants were sequenced to verify their accuracy. *S. pombe* genomic sequences and annotation from PomBase (Lock et al., 2019).

For serial dilution growth assays, cells were cultured in liquid YE at 32°C until mid-log phase, three 10-fold serial dilutions starting at 4 × 10^6^ cells/mL were made, 4μL of each dilution was spotted on YE plates, and cells were grown at the indicated temperatures for 2-4 d. For overexpression experiments, pREP1 plasmids containing *hhp1, hhp2*, and the corresponding mutants were transformed into a wildtype strain using electroporation and selected on minimal media containing thiamine (MAUT). Single colonies were grown to log phase in MAUT, then expression was induced by washing into minimal media without thiamine (MAU) and incubating at 32°C for 16-24h. Growth assays were performed as above, except all dilutions were in MAU, and cells were spotted on MAU and MAUT plates. Heat shock viability assay was performed as described previously (Plyte et al., 1996).

### Live cell imaging

Single colonies of pREP1 transformants were grown to log phase in MAUT, then expression was induced by washing into MAU and incubating at 32°C for 24h. Live-cell images of *S. pombe* cells were acquired using a Personal DeltaVision microscope system (Applied Precision) that includes an Olympus IX71 microscope, 60× NA 1.42 PlanApo and 100× NA 1.40 UPlanSApo objectives, a Photometrics CoolSnap HQ2 camera, and softWoRx imaging software.

### Protein alignment and structural modeling

The structural model of Hhp1 was generated using the Protein Homology/analogY Recognition Engine V 2.0 (Phyre2) (Kelley et al., 2015) as previously described (Elmore et al., 2018). Other structures were downloaded from the Protein Data Bank (rcsb.org; Berman et al., 2000), and all models were visualized in The PyMOL Molecular Graphics System, Version 1.8 Schrödinger, LLC. Primary sequence alignments were performed in COBALT (https://www.ncbi.nlm.nih.gov/tools/cobalt/cobalt.cgi) using reference sequences from UniProt (The Uniprot Consortium, 2019).

## Acknowledgements

We thank Rodrigo Guillen, Eric Zhang, Janel Beckley, Zachary Elmore, and Anna Feoktistova for helpful discussions and assistance with the CK1 project. We thank members of the Gould laboratory for critical comments on the manuscript. S.N.C. was supported by the Integrated Biological Systems Training in Oncology Program (T32-CA119925). This work was funded by R35-GM131799 to K.L.G.

