## Supplemental Material for "Autophosphorylation of the CK1 kinase domain regulates enzyme activity and function"

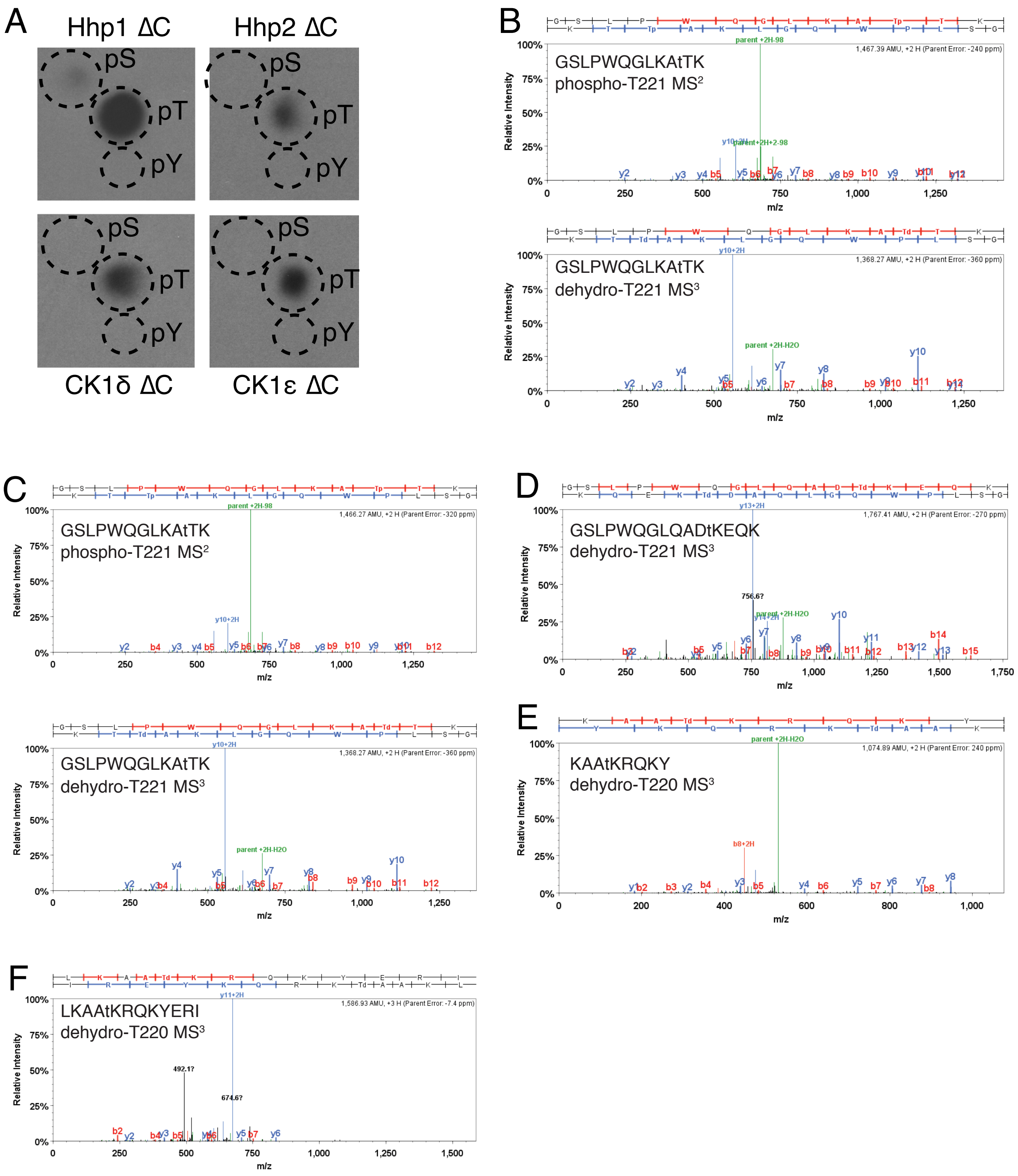


Supplemental Figure 1

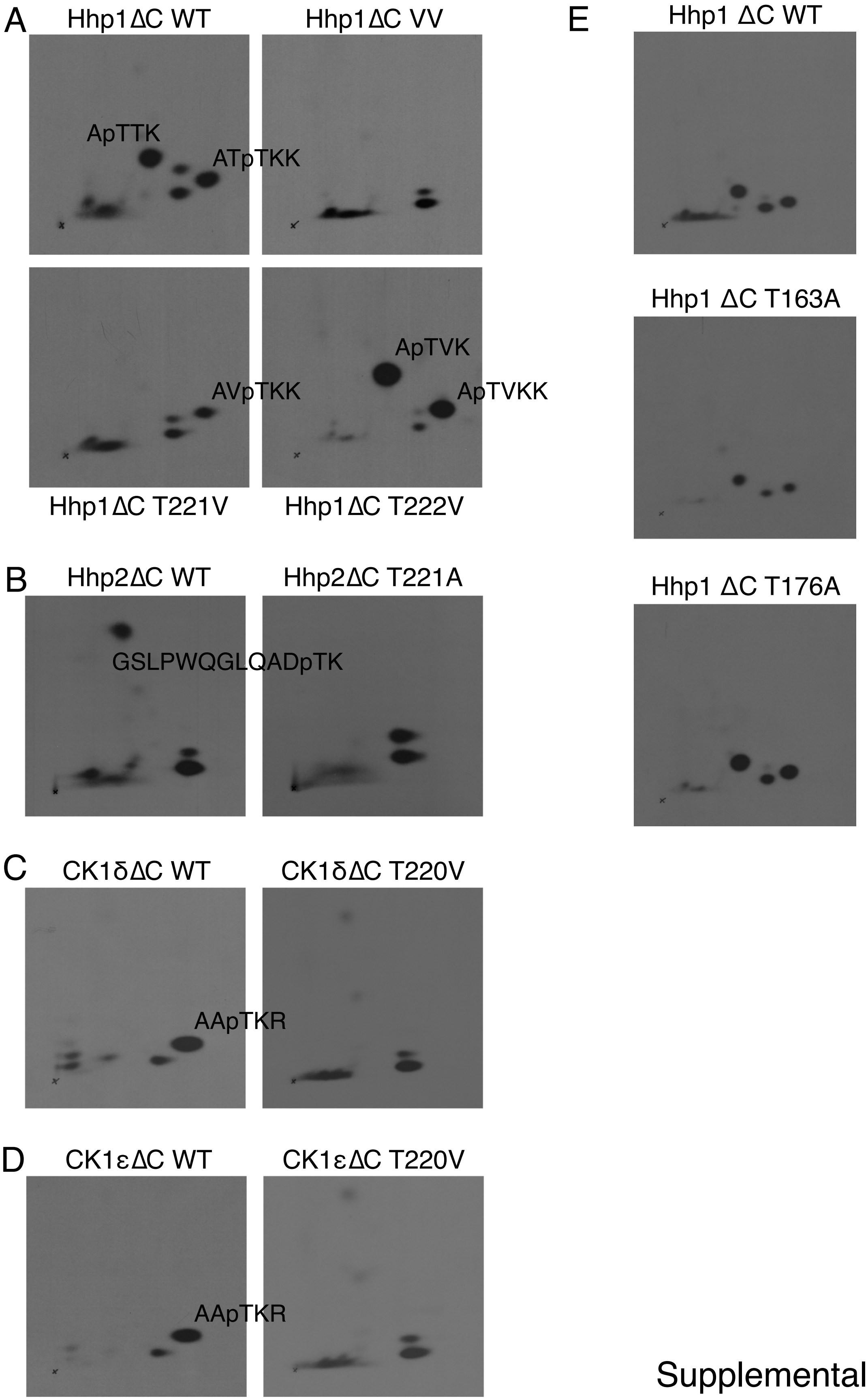


Supplemental Figure 2

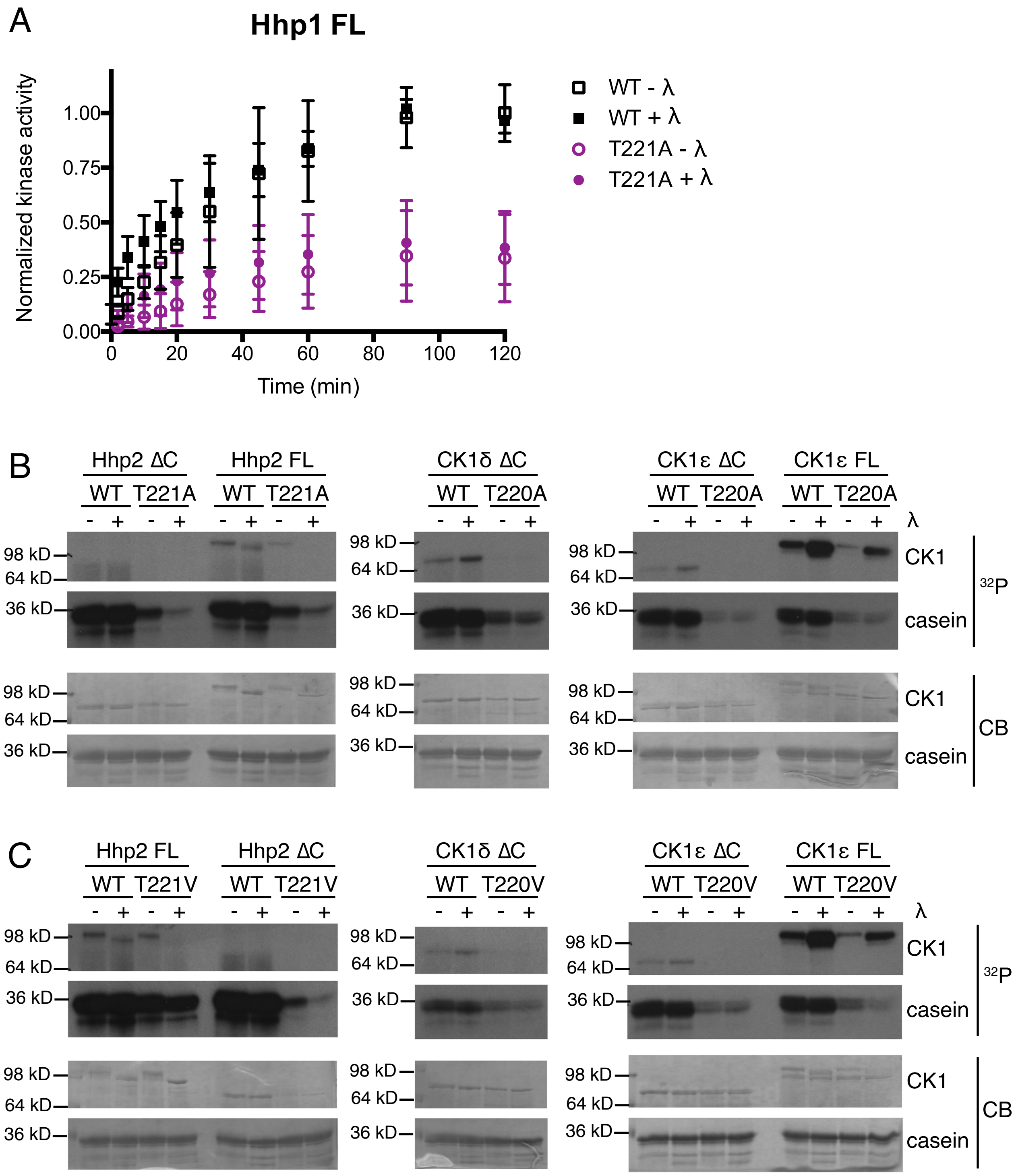


Supplemental Figure 3

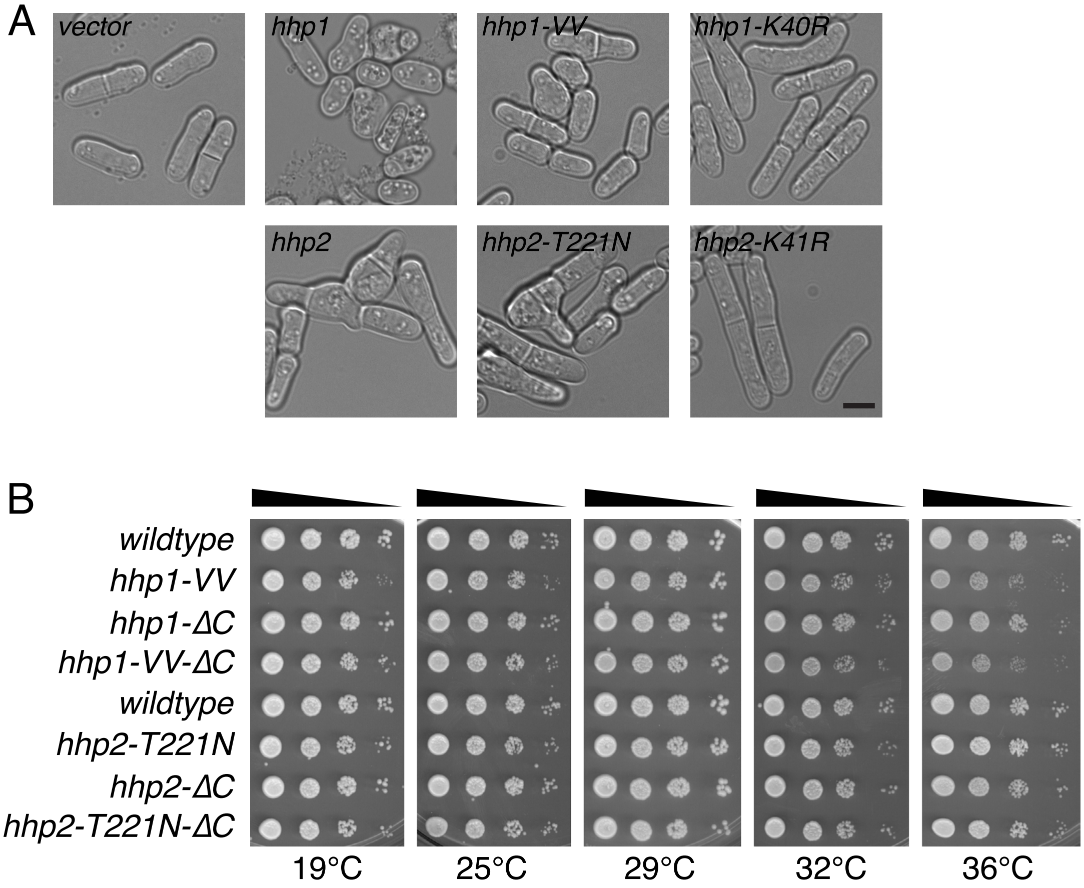


Supplemental Figure 4

**Supplemental Table 1: *S. pombe* strains used in this study.**

| Strain | Genotype | Reference |
|---|---|---|
| | Figure 5 | |
| 599 | *ura4-D18 leu1-32 h-* | Lab stock |
| 5106-2 | *hhp1-VV ura4-D18 leu1-32 h-* | This study |
| 13360-2 | *hhp1-ΔC:kan*$^{R}$ *ura4-D18 leu1-32 h-* | This study |
| 13389-2 | *hhp1-VV-ΔC:kan*$^{R}$ *ura4-D18 leu1-32 h-* | This study |
| 246 | *ura4-D18 leu1-32 ade6-M210 h-* | Lab stock |
| 5202-2 | *hhp2-T221N ura4-D18 leu1-32 ade6-M210 h+* | This study |
| 13414-2 | *hhp2-ΔC:nat*$^{R}$ *ura4-D18 leu1-32 h-* | This study |
| 13415-2 | *hhp2-T221N-ΔC:nat*$^{R}$ *ura4-D18 leu1-32 ade6-M210 h+* | This study |
| 6415 | *hhp1::ura4*$^{+}$ *ura4-D18 leu1-32 h-* | Bimbo et al. 2005 |
| | Figure S1 | |
| 14161 | *hhp1-HA*$_{3}$*-TAP:kan*$^{R}$ *ura4-D18 leu1-32 ade6-M210 h-* | Johnson et al. 2013 |
| | Figure S4 | |
| 599 | *ura4-D18 leu1-32 h-* | Lab stock |
| 5106-2 | *hhp1-VV ura4-D18 leu1-32 h-* | This study |
| 13360-2 | *hhp1-ΔC:kan*$^{R}$ *ura4-D18 leu1-32 h-* | This study |
| 13389-2 | *hhp1-VV-ΔC:kan*$^{R}$ *ura4-D18 leu1-32 h-* | This study |
| 246 | *ura4-D18 leu1-32 ade6-M210 h-* | Lab stock |
| 5202-2 | *hhp2-T221N ura4-D18 leu1-32 ade6-M210 h+* | This study |
| 13414-2 | *hhp2-ΔC:nat*$^{R}$ *ura4-D18 leu1-32 h-* | This study |
| 13415-2 | *hhp2-T221N-ΔC:nat*$^{R}$ *ura4-D18 leu1-32 ade6-M210 h+* | This study |

**Supplemental Figure 1: Identification of threonine autophosphorylation in the CK1 kinase domain.** (A) Truncated, MBP-tagged CK1 was treated with lambda phosphatase, then incubated with γ-[$^{32}$P]-ATP at 30°C for 30 min. Proteins were subjected to partial acid hydrolysis and separated by two-dimensional thin-layer electrophoresis. Phosphorylated amino acids were detected by autoradiography. Kinase domain autophosphorylation occurs on threonine. (B-F) CK1 enzymes were analyzed for post-translational modifications by mass spectrometry. Representative spectra of kinase domain autophosphorylation sites are shown. Hhp1-TAP affinity purified from *S. pombe* cells (B). Recombinant MBP-Hhp1 (C), MBP-Hhp2 (D), and MBP-CK1δ (E). CK1ε-MAP affinity purified from HEK293 cells (F).

**Supplemental Figure 2: Phosphopeptide mapping of kinase domain threonine mutants.** Truncated, MBP-tagged CK1 was treated with lambda phosphatase, then incubated with γ-[$^{32}$P]-ATP at 30°C for 30 min. Proteins were digested with trypsin, and peptides were separated by thin-layer electrophoresis and chromatography. Phosphopeptides were detected by autoradiography. (A) Maps of MBP-Hhp1ΔC WT, T221V, T222V, and VV show that both T221 and T222 can be autophosphorylated in vitro. Signals for the homologous phosphopeptides in MBP-Hhp2ΔC (B), MBP-CK1δΔC (C), and MBP-CK1εΔC (D) are abolished by mutating the conserved kinase domain threonine. (E) Threonines within the L-9D loop of Hhp1 are not autophosphorylated. The L-9D loop of CK1 is homologous to the activation loop (T-loop) of other RD kinases; however, CK1 does not require T-loop phosphorylation to be active. T163 and T176 are homologous to T-loop phosphorylation sites in other kinases, but they are not autophosphorylated by Hhp1, as shown by phosphopeptide maps of MBP-Hhp1ΔC T163A and T176A.

**Supplemental Figure 3: The conserved region surrounding the kinase domain threonine is sensitive to phosphoablating mutations.** (A) MBP-Hhp1 WT or T221A were treated with lambda phosphatase or mock treated, then incubated with casein and γ-[$^{32}$P]-ATP at 30°C. Reactions were quenched at time points from 0-120 min, and casein phosphorylation was measured on a phosphorimager. Hhp1 T221A has reduced activity, even though phosphorylation on T221 is inhibitory. (B-C) MBP-tagged CK1 enzymes were treated with lambda phosphatase or mock treated, then incubated with casein and γ-[$^{32}$P]-ATP at 30°C for 1 h. Phosphorylated proteins were visualized by autoradiography ($^{32}$P) and total protein by Coomassie (CB). Threonine to alanine mutations (B) also disrupt activity in Hhp2, CK1δ, and CK1ε. Threonine to valine mutations (C) are tolerated by Hhp1 but disrupt activity in Hhp2, CK1δ, and CK1ε.

**Supplemental Figure 4: High levels of unregulated CK1 cause defects in *S. pombe* growth and morphology but low levels are buffered.** (A) A plasmid encoding the indicated *hhp1* and *hhp2* genes under the control of the *nmt1* promoter was transfected into cells, and expression was induced in thiamine-free media for 24 h. Live cells were imaged at 25°C. Scale bar = 5μm. Defects in cell polarity and cytokinesis are evident with overexpression of the wildtype or autophosphorylation mutant alleles, while the kinase-dead alleles phenocopy the deletions. (B) Serial 10-fold dilutions of the indicated strains were grown at the indicated temperatures. All genes were expressed from the endogenous loci.
